# Monoclonal antibodies against human coronavirus NL63 spike

**DOI:** 10.1101/2023.03.12.532265

**Authors:** Meghan Conlan, Hyon-Xhi Tan, Robyn Esterbauer, Andrew Kelly, Stephen J Kent, Adam K Wheatley, Wen Shi Lee

## Abstract

The COVID-19 pandemic has illustrated the potential for monoclonal antibody therapeutics as prophylactic and therapeutic agents against pandemic viruses. No such therapeutics currently exist for other human coronaviruses. NL63 is a human alphacoronavirus that typically causes the common cold and uses the same receptor, ACE2, as the highly pathogenic SARS-CoV and SARS-CoV-2 pandemic viruses. In a cohort of healthy adults, we characterised humoral responses against the NL63 spike protein. While NL63 spike and receptor binding domain-specific binding antibodies and neutralisation activity could be detected in plasma of all subjects, memory B cells against NL63 spike were variable and relatively low in frequency compared to that against SARS-CoV-2 spike. From these donors, we isolated a panel of antibodies against NL63 spike and characterised their neutralising potential. We identified potent neutralising antibodies that recognised the receptor binding domain (RBD) and other non-RBD epitopes within spike.

## Introduction

Coronaviruses pose a significant zoonotic threat, as illustrated by the three major spillover events into humans in the past two decades by SARS-CoV, MERS-CoV and SARS-CoV-2. In addition to these highly pathogenic coronaviruses, there are four endemic human coronaviruses (HCoV-HKU1, -OC43, -NL63, -229E) that cause mild upper respiratory tract infections. HCoV infection and seroconversion typically happens in early childhood (1), with periodic re-infections throughout life, often occurring within 9 to 12 months following the last infection (2). The seven coronaviruses that infect humans can be divided into the α genus (NL63, 229E) and β genus (HKU1, OC43, SARS-CoV, MERS-CoV, SARS-CoV-2). Interestingly, NL63 binds to the same receptor (ACE2) as the highly pathogenic SARS-CoV and SARS-CoV-2 pandemic viruses (3), though NL63 usually only causes the common cold. NL63 was first discovered in 2004 within a 7-month-old baby in the Netherlands (4) and is thought to have been circulating in humans for many centuries with a bat origin (5). Importantly, a subgenotype associated with severe lower respiratory tract infection was reported in China in 2018 (6), indicating that ongoing viral evolution can result in more pathogenic variants.

The spike (S) protein of coronaviruses mediates entry into host cells and is the major target for vaccines and therapeutics. S consists of two subunits: S1, which contains the receptor binding domain (RBD) and N-terminal domain (NTD); and S2, which contains the fusion machinery. RBD represents the major neutralising target for SARS-CoV-2 (7), though neutralising antibodies against the NTD and S2 domains have been reported (8–10). Whether this neutralisation hierarchy is retained in NL63 remains unknown. Notably, there are significant structural differences between the S proteins of α- and β-HCoVs, with RBD being locked in a “down” conformation in the prefusion state and the duplication of NTD both canonical features of α-HCoVs (11–13). Interestingly, NL63 S is unable to readily bind to ACE2 in the prefusion state as the receptor binding residues within RBD are buried through interactions with NTD of the same protomer (13). It is thought that NTD interacting with heparan sulfate proteoglycans on the cell surface is required to allow RBD to subsequently bind ACE2. While major advances have been made in developing vaccines and antiviral therapeutics against SARS-CoV-2 (and related pathogenic β-HCoVs), there remain no vaccines or therapeutics available for endemic HCoVs. Here we characterised the humoral response to NL63 S and isolated a panel of NL63 S-specific monoclonal antibodies (mAbs).

## Results

### Humoral response to NL63 S

We recruited a cohort of 14 healthy adults and assessed antibody and B cell responses to NL63 S. We detected plasma IgG against both NL63 S and RBD in all subjects, with endpoint titres ranging from 1:678 to 1:6887 for S and 1:2506 to 1:8299 for RBD (Fig 1A). Using a live-virus neutralisation assay, we next measured plasma neutralisation titres (ID_50_) against NL63, which ranged from 1:29 to 1:745 (Fig 1A). Interestingly, neither Spike nor RBD IgG titres correlated with neutralisation titres (Fig S1). Next, we used fluorescently labelled spike probes to detect memory B cells (CD19+IgD-IgG+) specific for NL63 S, and SARS-CoV-2 S as a positive control since all subjects received at least 2 doses of a COVID vaccine (Fig 1B, gating in Fig S2). NL63 S-specific memory B cells were significantly lower than SARS-CoV-2 S-specific memory B cells, likely a result of being more recently vaccinated against COVID than being infected with NL63 (Fig 1C).

**Figure 1.**
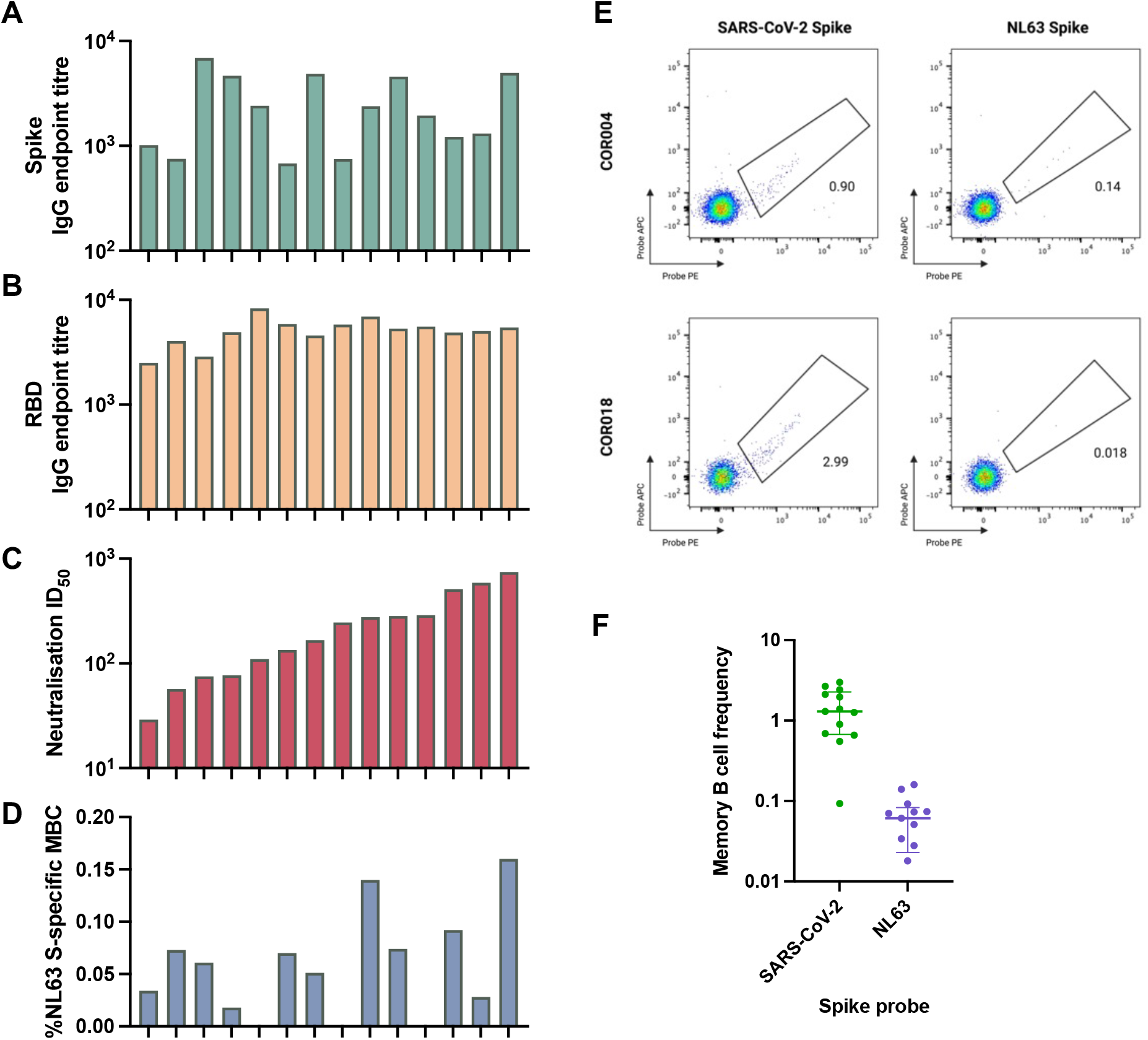
Humoral responses against NL63. S. Plasma samples from a cohort of healthy adults (N=14) were screened by ELISA for IgG antibodies against **(A)** NL63 S and **(B)** NL63 RBD. **(C)** NL63 neutralisation activity in plasma was assessed using a live virus neutralisation assay. **(D)** Frequencies of NL63 spike-specific memory B cells as a proportion of CD19+IgD-IgG+ B cells in PBMCs. **(E)** Representative flow cytometry plots of SARS-CoV-2 and NL63 S-specific memory B cells from two subjects stained with recombinant S proteins fluorescently labelled with APC or PE. **(F)** Frequencies of SARS-CoV-2 vs NL63 S-specific memory B cells as a proportion of CD19+IgD-IgG+ B cells in PBMCs (N=14).

### Monoclonal antibodies against NL63 S

Having identified the two subjects with highest memory B cell frequencies against NL63 S and decent plasma neutralisation titres (COR344 and COR004), we next single cell sorted NL63 S-specific memory B cells and recovered recombined immunoglobulin gene sequences using multiplex RT-PCR. A total of 52 heavy chain immunoglobulins and 91 light chains were recovered, with 13 paired heavy and light chains selected for expression as mAbs (denoted with prefix NL) in mammalian cell culture.

Seven of the 13 expressed human mAbs bound S and 4 of these were specific for RBD (Fig 2A, 2B, 2D). The S-binding mAbs were further screened for neutralising activity against NL63 (Fig 2C, 2D). Three potently neutralising mAbs were identified with IC_50_ ranging from 3.1 ng/ml to 67.5 ng/ml, with two mAbs binding to RBD (NL_2D04 and NL_3D09) and one mAb recognising a non-RBD epitope within S (NL_2B02). The RBD-specific mAb NL_3D09 only partially neutralised NL63 with a low IC_50_ of 3393 ng/ml.

**Figure 2.**
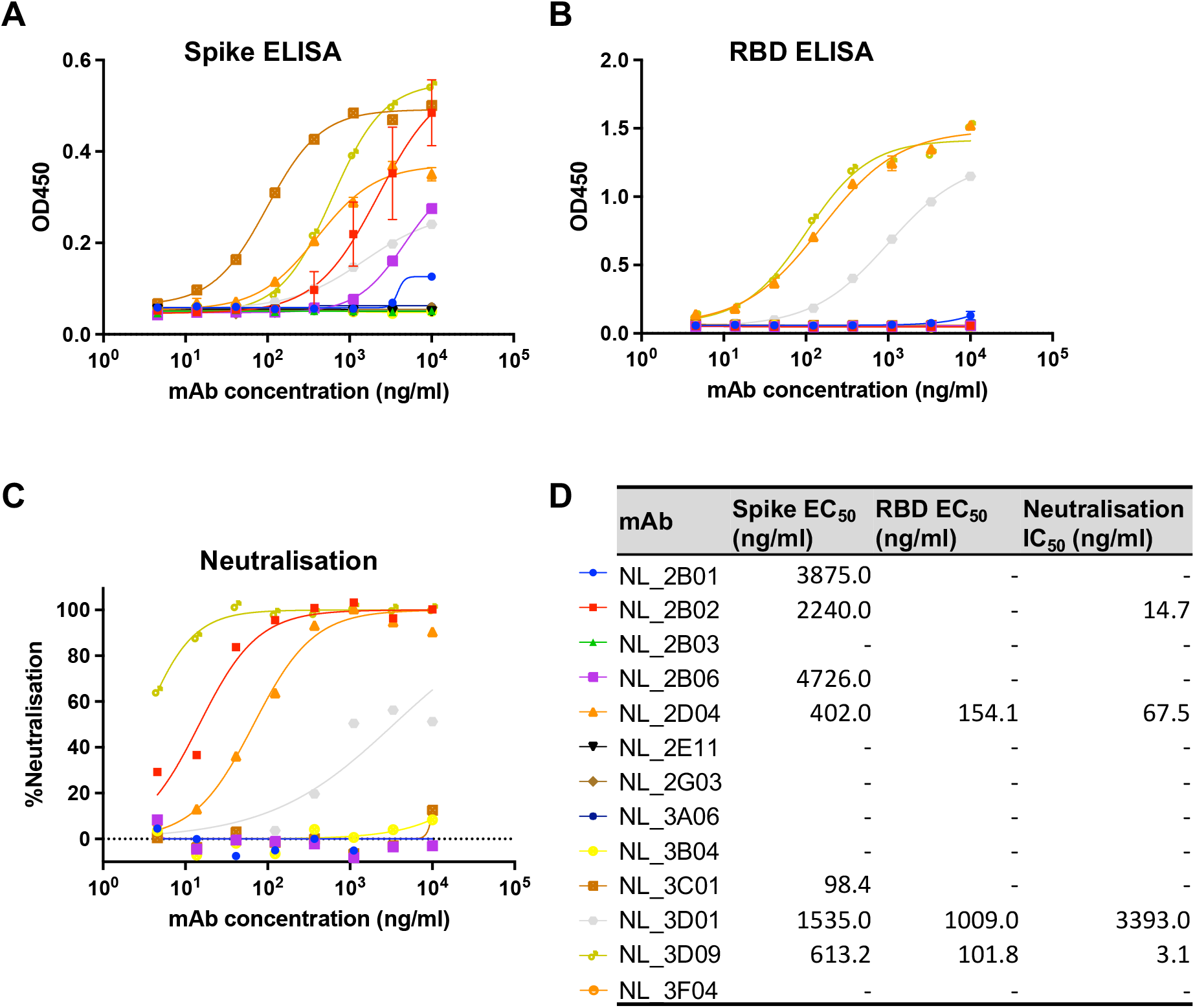
Monoclonal antibodies against NL63. **(A)** 14 mAbs were tested for reactivity against NL63 S by ELISA. 8 of the S-binding mAbs were tested further for reactivity against **(B)** NL63 RBD by ELISA and **(C)** neutralisation activity against NL63. **(D)** Table summarising the binding EC50 and neutralisation IC50 of the NL63 mAbs.

## Discussion

We characterised a panel of monoclonal antibodies against HCoV-NL63 spike. Four of these were neutralising, with three recognising RBD and one binding to a non-RBD epitope within S. While the RBD represents the predominant neutralisation target for SARS-CoV-2, whether this remains the case for NL63 remains unknown. Antigenic supersites within the NTD that harbour neutralising epitopes have been reported for SARS-CoV-2 (8, 10), OC43 (14), and 229E (15). A previous study found the NTD of 229E to be antigenically dominant, with mAbs displaying cross-reactivity with NL63, though cross-neutralising activity was not assessed (15). This site has the potential to be a pan-α-HCoV epitope for vaccine development if neutralisation activity is conserved across α-HCoVs. In addition to the NTD, several pan-β-HCoV and pan-CoV mAbs directed against conserved sites within the S2 domain have been reported (16–19). One such mAb, 76E1, recognises a highly conserved epitope between the S2’ site and fusion peptide and could neutralise 6 human coronaviruses, with IC_50_ values ranging from 0.4-4.8μg/ml (16). While the development of pan-CoV mAbs sounds promising, neutralising mAbs directed against the S2 domain tend to be less potent as they do not block receptor engagement and often do not bind S in the prefusion state. Instead, S2 mAbs neutralise virions by blocking membrane fusion steps that occur following receptor engagement. Therefore, the lower potency and resulting need for higher dosage may limit their prophylactic and therapeutic potential.

Interestingly, our study found that neither total S- or RBD-binding IgG antibodies correlated with NL63 neutralising activity. Given the duplication of NTD within α-HCoVs, whether NTD is the predominant target of neutralisation for NL63 remains to be determined. Two of the most potently neutralising NL63 mAbs we isolated (NL_2B02 and NL_3D09) have IC_50_ values similar to our most potent SARS-CoV-2 mAbs (20), indicating their potential for prophylactic or therapeutic development. Further characterisation of the epitopes recognised by these mAbs and their *in vivo* protective efficacies are ongoing.

## Materials and methods

### Human subjects and ethics

A cohort of 14 healthy adults was recruited to donate blood via venepuncture into 9ml blood collection tubes with sodium heparin anticoagulant. Plasma was collected and stored at −80°C, and PBMCs were isolated via Ficoll Paque separation, cryopreserved in 10% DMSO/FCS and stored in liquid nitrogen. The study protocols were approved by the University of Melbourne Human Research Ethics Committee (#2056689) and all associated procedures were carried out in accordance with the approved guidelines. All participants provided written informed consent in accordance with the Declaration of Helsinki.

### Expression of coronavirus antigens

A set of trimeric, pre-fusion stabilised coronavirus S proteins (NL63, SARS-CoV-2) were generated for serological and flow cytometric assays using techniques previously described (21). Genes encoding the ectodomain of SARS-CoV-2 S (NC_045512; AA1-1209) with 6 proline stabilisation mutations and furin site removal (Hexapro (22)), NL63 S (DQ445911.1; AA1-1291) with 2 proline stabilisation mutations (S-2P), and NL63 RBD were cloned into mammalian expression vectors. Proteins were expressed in Expi293 or ExpiCHO cells (Thermo Fisher, Massachusetts USA) using manufacturer’s instructions and purified using Ni-NTA and size exclusion chromatography. Protein integrity was confirmed using SDS-PAGE.

### ELISA against NL63 S and RBD

Antibody binding to HCoV-NL63 S and RBD protein was tested by ELISA. 96-well Maxisorp plates (Thermo Fisher) were coated overnight at 4°C with 2μg/mL recombinant S and RBD. After blocking with 1% FCS in PBS, duplicate wells of 4-fold serially diluted plasma (starting from 1:100) or mAbs (starting from 10μg/ml) were added and incubated for two hours at room temperature. Bound antibody was detected using 1:20,000 dilution of HRP-conjugated anti-human IgG (Sigma) and plates developed using TMB substrate (Sigma), stopped using sulphuric acid and read at 450nm. Endpoint titres were calculated using Graphpad Prism as the reciprocal serum dilution giving signal 2× background using a fitted curve (4 parameter log regression).

### NL63 virus propagation and titration

HCoV-NL63 (Amsterdam-1) isolate was grown in LLC-MK2 cells in DMEM with 2% FCS and 1μg/ml TPCK trypsin at 35°C. Cell culture supernatants containing infectious virus were harvested on Day 6, clarified via centrifugation, filtered through a 0.45μM cellulose acetate filter and stored at −80°C. Infectivity of virus stocks was then determined by titration on HAT-24 cells (a clone of transduced HEK293T cells stably expressing human ACE2 and TMPRSS2 (23)). In a 96-well flat bottom plate, virus stocks were serially diluted five-fold (1:5-1:10,935) in DMEM with 5% FCS, added with 30,000 freshly trypsinised HAT-24 cells per well and incubated at 35°C. After 70 hours, 10μl of alamarBlue™ Cell Viability Reagent (ThermoFisher) was added into each well and incubated at 35°C for 1 hour. The reaction was then stopped with 1% SDS and read on a FLUOstar Omega plate reader (excitation wavelength 560nm, emission wavelength 590nm). The relative fluorescent units (RFU) measured were used to calculate %viability (‘sample’ ÷ ‘no virus control’ × 100), which was then plotted as a sigmoidal dose response curve on Graphpad Prism to obtain the virus dilution that induces 50% cell death (50% infectious dose; ID_50_). Each virus was titrated in quintuplicate in three independent experiments to obtain mean ID_50_ values.

### NL63 neutralisation assay with viability dye readout

In 96-well flat bottom plates, 3-fold dilutions of heat-inactivated plasma samples (1:20-1:43,740) or NL63 mAbs (10,000 – 4.6 ng/ml) were incubated in duplicate with NL63 virus at a final concentration of 2× ID_50_ at 35°C for 1 hour. Next, 30,000 freshly trypsinised HAT-24 cells in DMEM with 5% FCS were added and incubated at 35°C. ‘Cells only’ and ‘Virus+Cells’ controls were included to represent 0% and 100% infectivity respectively. After 70 hours, 10μl of alamarBlue™ Cell Viability Reagent (ThermoFisher) was added into each well and incubated at 37°C for 1 hour. The reaction was then stopped with 1% SDS and read on a FLUOstar Omega plate reader (excitation wavelength 560nm, emission wavelength 590nm). The relative fluorescent units (RFU) measured were used to calculate %neutralisation with the following formula: (‘Sample’ – ‘Virus+Cells’) ÷ (‘Cells only’ – ‘Virus+Cells’) × 100. IC_50_ values were determined using four-parameter non-linear regression in GraphPad Prism with curve fits constrained to have a minimum of 0% and maximum of 100% neutralisation.

### Flow cytometric detection of spike-reactive B cells

Biotinylated recombinant NL63 spike protein was conjugated to streptavidin-APC or - PE fluorophores. PBMCs were thawed and stained with Aqua viability dye (Thermo Fisher Scientific) and then surface stained with Spike probes, CD19 ECD (J3-119) (Beckman Coulter), IgA VioBlue (IS11-8E10) IgM BUV395 (G20-127), IgD PE-Cy7 (IA6-2), IgG BV786 (G18-145), CD21 BUV737 (B-ly4), CD27 BV605 (O323), streptavidin BV510 (BD Biosciences), CD14 BV510 (M5E2), CD3 BV510 (OKT3), CD8a BV510 (RPA-T8), CD16 BV510 (3G8), and CD10 BV510 (HI10a) (BioLegend). Cells were washed twice with PBS containing 1% FCS and fixed with 1% formaldehyde (Polysciences) and acquired on a BD LSR Fortessa using BD FACS Diva.

### Recovery of human monoclonal antibodies from NL63 spike-specific B cells

NL63 spike-specific B cells were identified within cryopreserved human PBMC using fluorescent spike probes as described above. Single antigen-specific class-switched B cells (S or RBD+, CD19+ IgD-IgG+) were sorted using a BD Aria II into 96-well plates, subject to cDNA generation and multiplex PCR and Sanger sequencing, as previously described (21, 24). Productive, recombined heavy (V-D-J) and light (V-J) chain immunoglobulin sequences were synthesised (Geneart) and cloned into human IgG1 expression vectors for recombinant production in Expi293 mammalian cell culture using transient transfection. After 4 – 5 days, IgG1 was purified from culture supernatants using Protein-A affinity chromatography.

## Acknowledgements

We thank the participants for the generous involvement and provision of samples. We thank Dr Mingyang Wang and Prof Kanta Subbarao for isolating and distributing HCoV-NL63 virus isolates. We thank Lauren Burmas, Thakshila Amarasena, Devaki Pilapitiya, Grace Gare and Lara Schwab for technical assistance with ELISAs, cloning, protein expression and blood processing. We acknowledge the Melbourne Cytometry Platform for provision of flow cytometry services.

## Funding

Australian National Health and Medical Research Council Investigator or Fellowship grants (HXT, SJK, AKW)

Melbourne Postdoctoral Fellowship (WSL)

## Competing interests

The authors declare no competing interests.

## Data and materials availability

All data are available in the main text or the supplementary materials.

## Supplementary material

**Figure S1.**
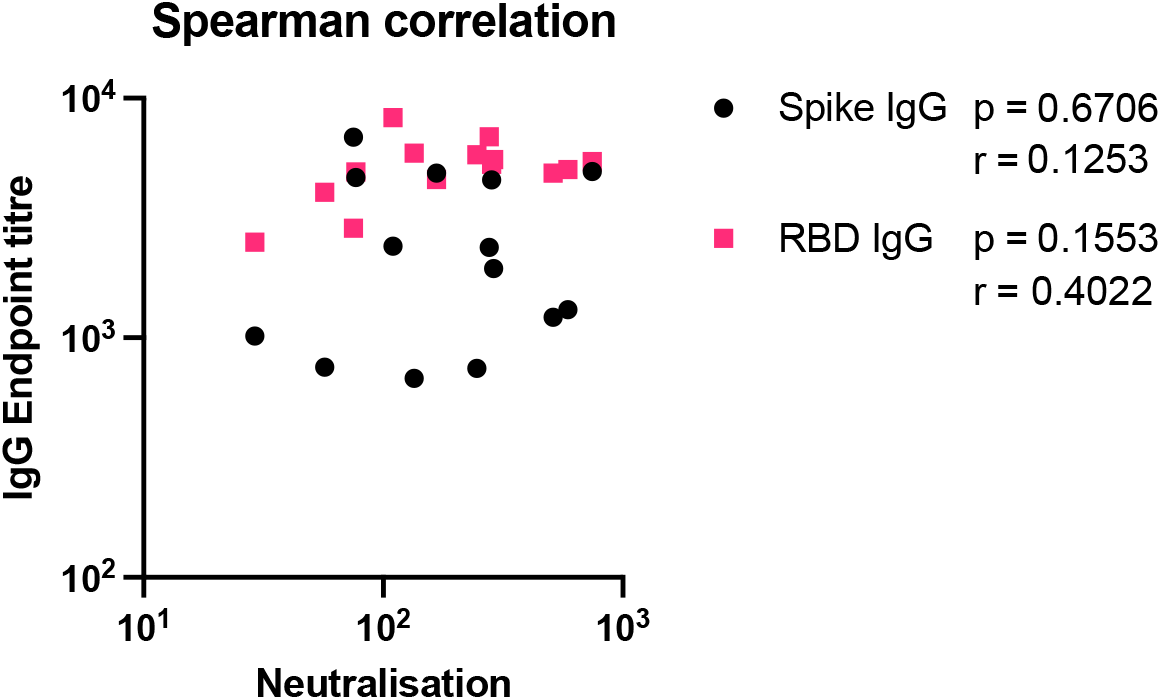
Spearman correlation of NL63 neutralisation activity against NL63 Spike and RBD IgG binding titres.

**Figure S2.**
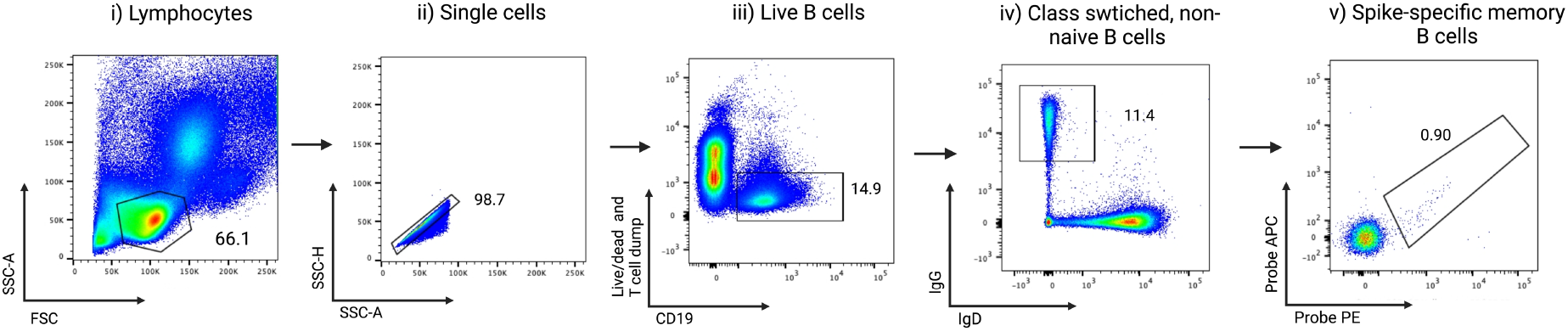
Flow cytometry gating strategy to identify SARS-CoV-2 and NL63 spike-specific memory B cells. (i) Lymphocytes were identified by FSC-A vs SSC-A gating, followed by (ii) doublet exclusion (SSC-A vs SSC-H), and (iii) gating on live CD19+ B cells. (iv) Class switched B cells were identified as CD19-IgG+ and (v) binding to recombinant SARS-CoV-2 or NL63 spike proteins conjugated to APC or PE was assessed.

## Notes

### Competing Interest Statement

The authors have declared no competing interest.

